# The role of brain oscillations in predicting the sensory consequences of your actions

**DOI:** 10.1101/065961

**Authors:** Liyu Cao, Gregor Thut, Joachim Gross

**Affiliations:** Institute of Neuroscience and Psychology, University of Glasgow, Glasgow, UK

**Keywords:** neural oscillations, auditory perception, sensory attenuation, prediction

## Abstract

Being able to predict self-generated sensory consequences is an important feature of normal brain functioning. In the auditory domain, self-generated sounds lead to smaller brain responses compared to externally generated sounds. Here we investigated the role of brain oscillations underlying this effect. With magnetoencephalography, we show that self-generated sounds are associated with increased pre-stimulus alpha power and decreased post-stimulus gamma power and alpha/beta phase locking in auditory cortex. All these oscillatory changes are correlated with changes in evoked responses. Furthermore, they correlate with each other across participants, supporting the idea that they constitute a neural information processing sequence for self-generated sounds, with pre-stimulus alpha power representing prediction and post-stimulus gamma power representing prediction error, which is further processed with post-stimulus alpha/beta phase resetting. Additional cross-trial analysis provides further support for the proposed sequence that might reflect a general mechanism for the prediction of self-generated sensory input.

## Introduction

In our interactions with the environment, action and perception are tightly linked. Voluntary motor actions typically lead to predictable sensory consequences. For example, knocking on a door results in a predictable sensory input to the auditory and somatosensory systems. It is well established that these self-generated sensory stimuli elicit smaller brain responses than externally generated stimuli (Blakemore, Wolpert, & Frith, 1998; Martikainen, Kaneko, & Hari, 2005; Schafer & Marcus, 1973) - a phenomenon known as sensory attenuation (SA). For example, an MEG study showed a reduced auditory M100 component when the sound was generated by participants pressing a button compared to when the sound was passively presented (Martikainen et al., 2005).

A forward model has been proposed to account for this effect (Blakemore, Frith, & Wolpert, 1999; Ramnani, 2006; Wolpert & Ghahramani, 2000). The model posits that along with a motor command, an efference copy (von Holst & Mittelstaedt, 1950) is sent that allows the computation of the predicted, imminent sensory consequences. The predicted sensory signal is then compared to the actual incoming sensory signal and results in a modulation of the brain responses depending on the match between the real and the predicted sensory signal (attenuated when matching). A detailed conceptual explanation can be derived from the predictive coding theory (Friston, 2005). In this framework, the evoked response is an expression of prediction error, which is the discrepancy between the predicted sensory consequence and the actual sensory input. Accurately predicted stimuli lead to smaller prediction errors, which is reflected in a decreased evoked response. In addition, it has been suggested that predictions and prediction errors are communicated along cortical hierarchies in distinct frequency bands. More specifically, recent evidence suggests that predictions are communicated along anatomical feedback connections via alpha/beta rhythms and prediction errors are communicated along feedforward connections via gamma rhythms (Bastos et al., 2015; Michalareas et al., 2016; Wang, 2010).

Here, we studied auditory sensory attenuation (SA) with MEG to test the hypothesis that predictions and prediction errors related to the internal forward model are reflected in neural oscillations. Neural oscillations in lower frequency bands are likely candidates for the implementation of SA as they are tightly linked to excitability changes in neural populations (Jensen & Mazaheri, 2010; Thut, Miniussi, & Gross, 2012; Weisz, Hartmann, Müller, Lorenz, & Obleser, 2011), and therefore may mediate gain control for the processing of incoming sensory information. A number of studies provide converging evidence that low frequency oscillations particularly in the 10 Hz range (alpha band) support active inhibition. An increase in alpha power is typically associated with a decrease in perceptual performance (Frey et al., 2014; Thut, Nietzel, Brandt, & Pascual-Leone, 2006; Van Dijk, Schoffelen, Oostenveld, & Jensen, 2008). Moreover, the phase of low frequency oscillations (including alpha) was also shown to modulate neural excitability, so that near-threshold stimuli are more likely to be perceived or neural responses to be enhanced if stimulus presentation is aligned to a certain phase of the ongoing oscillations (Arnal & Giraud, 2012; Busch, Dubois, & VanRullen, 2009; Lakatos, Chen, O'Connell, Mills, & Schroeder, 2007; Mathewson, Gratton, Fabiani, Beck, & Ro, 2009). We therefore hypothesized that pre-stimulus changes in low frequency oscillations may reflect a prediction process, which is generated by the forward model to implement the suppression of post-stimulus responses for SA. Indeed, some studies already provided evidence that pre-stimulus alpha power is higher in the sensory cortex when speech or visual stimuli are self-induced by movement (Müller, Leske, Hartmann, Szebenyi, & Weisz, 2014; Stenner, Bauer, Haggard, Heinze, & Dolan, 2014).

We further hypothesised that processes related to prediction error are reflected in gamma oscillations (Bauer, Stenner, Friston, & Dolan, 2014; Behroozmand et al., 2016). This is also in line with findings showing that gamma oscillations relay feedforward information (e.g., Michalareas et al. (2016)). In the context of SA, intracranial recordings from neurosurgical participants showed that gamma power (70-150 Hz) was suppressed in response to speech stimuli during speaking as compared to listening (Flinker et al., 2010). Thus reduced gamma power may indicate decreased prediction errors when the stimulus is predicted through the forward model during speaking compared to listening. Therefore, we predicted reduced gamma power for self-generated versus passively presented stimuli in the present study.

However, a decrease in post-stimulus gamma power does not seem to contribute to SA as measured with trial-averaged evoked responses (e.g., reflected in attenuated M100 component), where a low pass filter at around 40 Hz was applied in many studies (e.g., Baess, Horváth, Jacobsen, and Schröger (2011); Martikainen et al. (2005); Müller et al. (2014)). Overall, our understanding about the neural processes underlying SA at the level of single trials is still incomplete. A reduced amplitude of evoked responses after averaging across trials during SA could result from an amplitude reduction in single trials, an increased single trial phase jitter or a combination of both. Since sensory evoked responses are primarily reflected in an increase in the power and phase locking of theta oscillation, one may expect that a reduction of power and/or phase locking in the same frequency band contributes to SA.

In summary, our study addresses the following questions: 1) How is the pre-stimulus prediction of expected sensory consequences of an action reflected in the oscillatory activity of sensory brain areas? 2) How is the post-stimulus attenuation of evoked field responses in SA represented in the frequency domain (as a decrease of oscillatory power, phase locking or both)? 3) How do the pre- and post-stimulus changes in oscillatory activity underlying SA interact? To answer these questions, we conducted an MEG experiment using a well-established SA paradigm in the auditory domain, in which neural responses from self-generated and passive stimuli were compared (Baess et al., 2011; Schafer & Marcus, 1973). After confirming the existence of SA in auditory cortex, we performed time-frequency analysis for neural activations in auditory cortex to answer these questions.

## Methods

### Participants, Procedure and Recording

14 healthy, right-handed volunteers (6 males, mean age = 22.6, SD = 1.8) were recruited from a local participants’ pool. Participants gave written informed consent prior to the experiment and received monetary compensation after the experiment. The study was approved by the local ethics committee (Ethics Committee of College of Science and Engineering, University of Glasgow) and was conducted in accordance with the Declaration of Helsinki. A 248-magnetometers whole-head MEG system (MAGNES 3600 WH, 4-D Neuroimaging) was used for data recording with a sampling rate of 1,017Hz.

The stimulus was a pure tone (1000 Hz, 50 ms in duration, 90 dB sound pressure level) delivered through a plastic tube. There were four conditions (100 trials each). In the passive periodic condition, the auditory stimulus was controlled by the computer and was presented once every three seconds. The passive jittered condition was the same with the passive periodic condition except that the stimulus was presented with a jittered interval (between 2000 and 4000 ms, uniform distribution). In the active condition, the stimulus was presented immediately after an index finger lifting movement that the participants were asked to perform about once every three seconds without inner counting. The motor only condition was the same with the active condition except that no stimulus was presented after each movement. We used a light sensor (instead of a response box) to record the movements without noise associated with the finger movement. Every movement unblocked the beam from the light sensor (placed next to participant’s right index finger), which then generated a sound stimulus. Participants were asked to close their eyes during testing. Before the start of the experiment, participants received 50 trials of practice to familiarize themselves with the light sensor and the rate of finger movements. During this practice, they were asked to move the finger about once every three seconds without inner counting and they received visual feedback for their timing performance after each trial. No such feedback was provided in the real data collection. The four conditions were presented in a random order and participants were encouraged to take a break in between. The condition with jittered stimulus presentation served to analyze spontaneous fluctuations in preparedness to sounds (after having identified the oscillatory correlates in the active vs passive periodic comparisons). The movement only condition was not further analyzed here.

### Data analysis

Data analysis was performed with Matlab using FieldTrip toolbox (Oostenveld, Fries, Maris, & Schoffelen, 2011) and in-house codes in accord with current MEG guidelines (Gross et al., 2013). Trials with very short inter-trial intervals (less than 1500 ms) were discarded (8.1% of all trials). Then MEG signals were denoised using ft_denoise_pca and trials with artifacts were removed following visual inspection with ft_rejectvisual. Eye movement and heart artefacts were rejected using ICA. On average 93.6 (SD: 4.1, minimum: 85) and 94.0 (SD: 4.4, minimum: 86) trials remained after this step for the active and passive periodic condition, respectively.

### Evoked Responses

In sensor space analysis, MEG signals were low-pass filtered with 30 Hz cut-off frequency. Original magnetometer signals were converted to planar gradient representation. Three sensors from each hemisphere that were predominantly responding at the latency of the M100 component (95-120 ms post-stimulus) in the passive periodic condition were selected for analysis. Event related fields aligned to the onset of the sound were computed for each condition with baseline (−500 to −100 ms) correction. The M100 component was statistically compared between conditions using a paired t-test with a Monte Carlo randomization with 1000 permutations.

### Source Localization

T1-weighted structural magnetic resonance images (MRIs) of each participant were co-registered to the MEG coordinate system using a semi-automatic procedure. Anatomical landmarks (nasion, left and right pre-auricular points) were manually identified in the individual's MRI. Initial alignment of both coordinate systems was based on these three points. Subsequently, numerical optimization was achieved by using the ICP algorithm (Besl & McKay, 1992).

Individual head models were created from anatomical MRIs using segmentation routines in FieldTrip/SPM5. Leadfield computation was based on a single shell volume conductor model (Nolte, 2003) using a 10 mm grid defined on the template (MNI) brain. The template grid was transformed into individual head space by linear spatial transformation.

The localization of the auditory evoked component was based on the eLoreta algorithm as implemented in Fieldtrip (http://www.fieldtriptoolbox.org/). All other analyses used LCMV filters computed based on a covariance matrix from −500 ms to 500ms with a regularisation of 7% of the mean across eigenvalues of the covariance matrix.

All further analyses were based on a representative voxel from the right primary auditory cortex that was anatomically defined and showed clear reconstructed evoked responses. We focused the analysis on the right auditory cortex because activity estimates for left auditory voxels can be contaminated by activity from the left primary motor cortex related to the movement of the right hand finger. For the selected voxel, we computed an LCMV filter along the orientation of maximal power (across all experimental conditions) and extracted the single-trial time series separately for each experimental condition.

### Time-frequency Analysis

For the selected voxel, we subjected the time-series to time-frequency analysis, separately for each participant and experimental condition. We performed time-frequency analysis with a temporal resolution of 10 ms and a spectral resolution of 1 Hz on 500 ms (200 ms) long sliding windows for frequencies below (above) 40 Hz. For frequencies below 40 Hz, a Hanning taper was applied before fourier transformation. For frequencies above 40 Hz, a Hanning taper was applied for phase estimation and a multi-taper approach was used for power estimation with a smoothing of 10 Hz.

To test for differences between conditions, individual time-frequency maps were subjected to dependent-sample t-test (active vs passive periodic). Oscillatory power was log transformed with reference to the mean power from −700 to 700 ms. The null distribution was estimated using 1000 randomizations and multiple comparison correction was performed using the cluster method (Maris & Oostenveld, 2007). Only significant results (p < 0.05, cluster correction) are reported.

### Correlation Analysis

We used Spearman correlation for all correlations across participants implemented in the robust correlation toolbox (Pernet, Wilcox, & Rousselet, 2012). First, SA was used for correlation with the pre-stimulus alpha power increase, post-stimulus alpha phase locking decrease, and post-stimulus gamma power decrease. SA was calculated by taking the relative change of evoked responses (70-160 ms) between the active and passive periodic condition using the voxel reconstructed time series data (i.e., the amplitude difference in evoked responses between the active and passive periodic condition divided by the amplitude of evoked responses in the passive periodic condition). We used the voxel with strongest M100 activation in right auditory cortex and used t-test to test if SA is significantly different from 0. Other components used for correlation analysis were derived from clusters showing significant differences between active and passive periodic conditions (see results). The pre-stimulus alpha power increase (10 Hz, −400 to 0 ms), post-stimulus alpha phase locking decrease (9-10 Hz, 0-150 ms), and post-stimulus gamma power decrease (high gamma: 85-104 Hz, 90-120 ms; low gamma: 57-62 Hz, 30-80 ms) refer to relative changes that were calculated in the same way as SA. Second, correlation analysis was performed among the oscillatory changes between conditions: the pre-stimulus alpha power increase (9-11 Hz, −400 to −60 ms) and the post-stimulus gamma power decrease (93-106 Hz, 90-120 ms); the post-stimulus alpha phase locking decrease (9-11 Hz, 0-150 ms) and the post-stimulus gamma power decrease (85-104 Hz, 80-110 ms).

### Analysis of Passive Jittered Condition Data

This part of analysis aims at corroborating the existence of an oscillatory neural information processing sequence (see Figure 5d) from the above between-condition comparisons by testing them in a single condition setting. In the passive jittered condition, single trial pre-stimulus alpha power (8-12 Hz, −300 to 0 ms) was extracted per participant and correlated with the single trial gamma power (absolute baseline correction; baseline: −300 to 0 ms). Pearson correlation coefficients were Fisher z transformed before being subjected to t-tests against 0. For analyzing the relationship between post-stimulus gamma power and post-stimulus alpha/beta phase, we computed the phase deviation as the absolute angular difference of a single trial phase to the mean phase across trials. Then we used the phase deviation in the time-frequency window that showed a significant difference between conditions (Figure 2b; 12 to 14 Hz, 70 to 160 ms) to correlate with single trial gamma power (with circ_corrcl from CircStat toolbox; Berens (2009)). The correlation coefficients within the first 100 ms after the stimulus onset were statistically compared to mean correlation coefficients in a baseline period (-300 ms to 0; paired t-test) after Fisher z transformation. Next, we correlated the gamma power (99-106 Hz, 0-40 ms; Figure 5b) from the time window where significant correlations were found with the alpha/beta (12-14 Hz) phase deviation over time (from −750 ms to 750 ms) to reveal the temporal relationship between them. Cluster correction was applied to all the multiple comparisons.

## Results

### Replication of sensory attenuation in auditory cortex

We replicated the typical sensory attenuation effect on auditory evoked fields (SA). There was a significant decrease in the amplitude of sound evoked M100 component in the active as compared to the passive periodic condition (left sensors: t(13) = −3.67, p < 0.01; right sensors: t(13) = −3.99, *p* < 0.01) (Figure 1b). Source localization analysis demonstrated the maximum SA effect in the auditory cortex, confirming a significant reduction of primary auditory cortex response amplitude for self-initiated sounds compared to external sounds (Figure 1c). No significant differences were found in M100 amplitudes between the two passive conditions (left sensors: t(13) = 0.22, *p* = 0.43; right sensors: t(13) = 0.33, *p* = 0.37).

**Figure 1.**
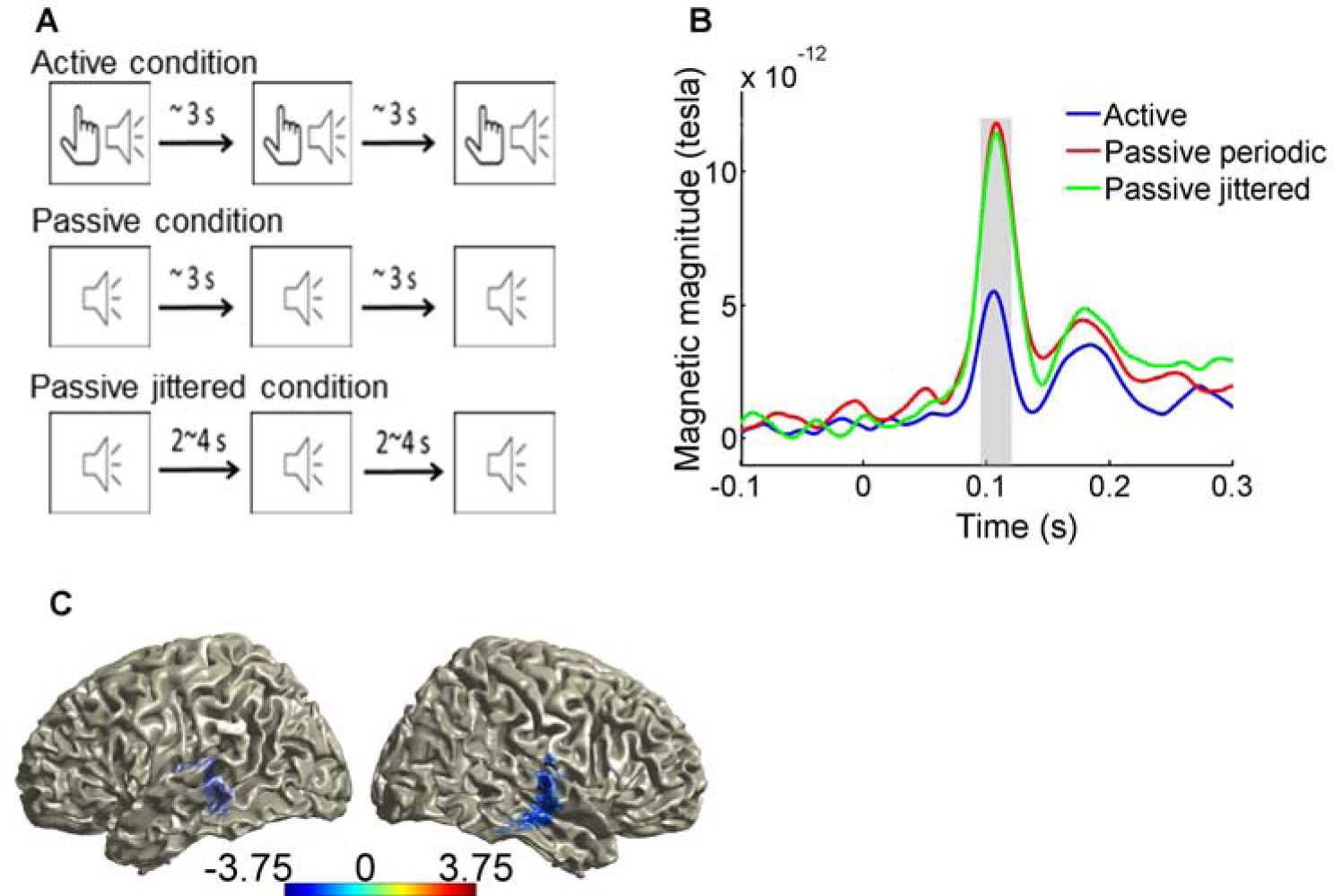
Replication of sensory attenuation effect. (A) schematic show of the active and passive periodic condition. (B) M100 component (the shaded area) is significantly lower in the active compared to the passive periodic condition (p < 0.01). (C) SA effect is localized in auditory area using eLoreta.

### Neural activity preceding sensory attenuation in auditory cortex

As introduced above, we hypothesised that in the SA auditory evoked responses are modulated by anticipatory prediction mechanisms through the forward model, possibly reflected in the alpha-band oscillation. Therefore, we tested for differences in oscillations between the active and passive periodic condition in the pre-stimulus period.

This analysis revealed differences in the state of auditory cortex between both conditions prior to the presentation of the stimulus. A significant increase in alpha band power (∼10 Hz) was found in the active as compared to the passive periodic condition starting around 400 ms before stimulus onset (Figure 2a). Testing for a relationship between this pre-stimulus alpha power increase and the magnitude of SA across participants revealed a significant correlation: increased alpha power was associated with increased SA (Spearman’s rho = −0.74, *p* = 0.003, 95% CI = [−0.92 −0.33]; Figure 3a, see also supplementary material for robust correlation results). Note that no significant changes in phase locking were found in the pre-stimulus time window (Figure 2b).

**Figure 2.**
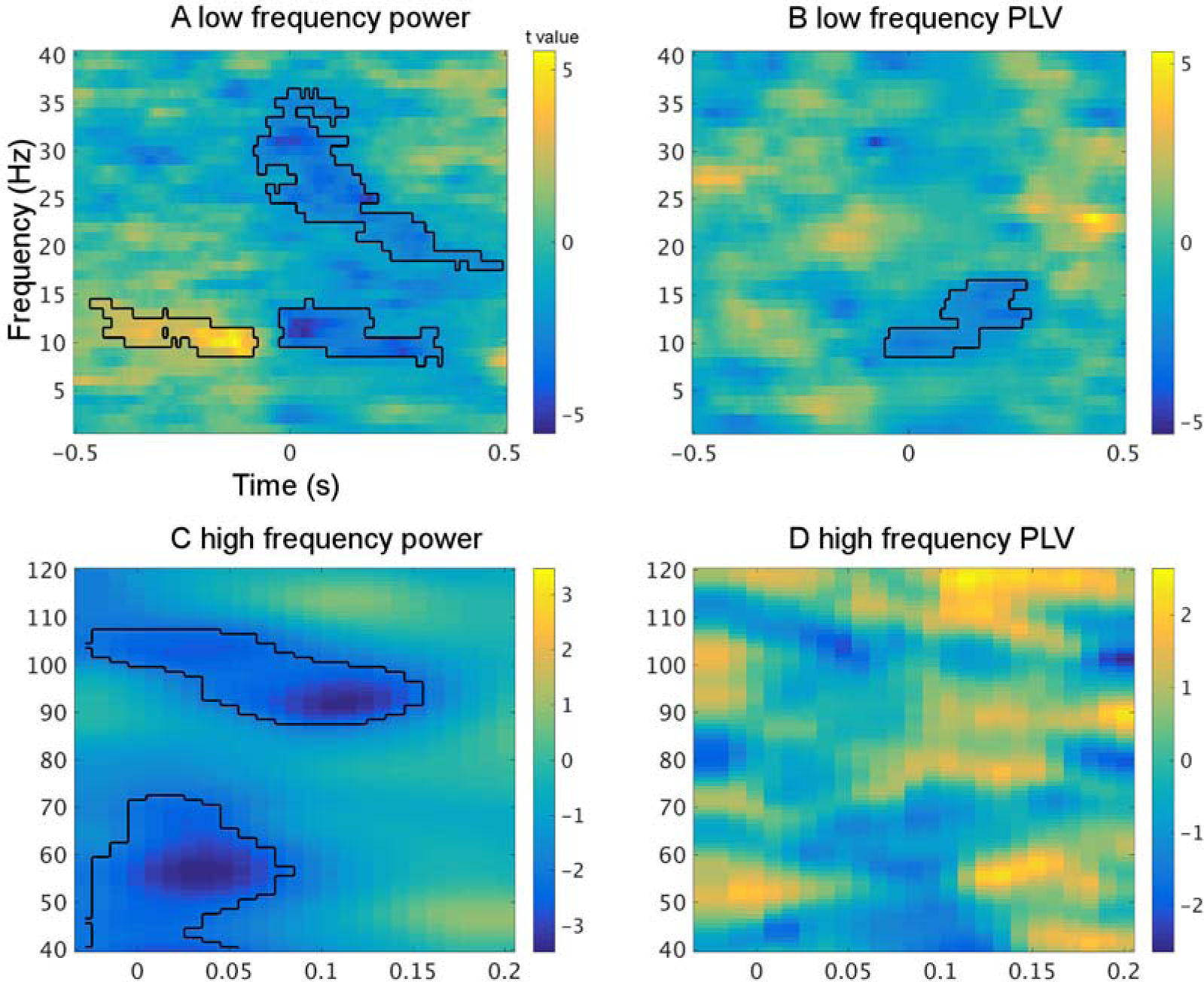
Power and phase locking value (PLV) comparisons between the active and passive periodic condition. In the pre-stimulus time window, a clear alpha power increase is shown (panel A). In the post-stimulus time window, broadband power decreases coincide with sensory attenuation from the evoked fields analysis (panel A and C). Of particular interest is the post-stimulus gamma power decrease. But there are no changes to theta band oscillations. Post-stimulus phase locking is decreased in the alpha/beta range (panel B). No difference is found in the gamma range phase locking (panel D).

### Neural representation of sensory attenuation in auditory cortex

To examine how SA is expressed in single-trials across different frequencies of neural activity, we focused next on frequency-specific activity in the post-stimulus time window that overlaps with the M100 component and statistically compared the oscillatory power and inter-trial phase locking between the active and passive periodic condition.

Time-frequency analysis revealed a significant decrease of broadband power at frequencies in the alpha/ low beta (9-15) and higher beta band (20-35 Hz) (Figure 2a) as well as in the gamma band (40-70 Hz and 90-110 Hz, Figure 2c) for the active as compared to the passive periodic condition. Time-frequency analysis of the passive jittered condition revealed that the evoked component was most strongly represented in the theta frequency band (see supplementary figure S1). Interestingly, differences between conditions only occurred at higher frequencies. These broadband changes overlapped in time with the SA effect. In parallel, phase locking to stimulus onset was significantly reduced in a limited frequency band, spanning the alpha/low beta frequency (9-15 Hz) in the same time window, for the active as compared to the passive periodic condition (Figure 2b). No significant differences were found in gamma band phase locking (Figure 2d).

When examining the relationship between these post-stimulus events and SA, we found the SA effect ( t(13) = −2.87, *p* = 0.01 in the voxel of peak oscillatory changes) to significantly correlate with three post-stimulus oscillatory events namely alpha phase locking decrease (Spearman’s rho = 0.63, *p* = 0.02, 95% CI = [0.05 0.94], Figure 3b), high gamma power decrease (Spearman’s rho = 0.69, *p* = 0.006, 95% CI = [0.29 0.90], Figure 3c) and low gamma power decrease (Spearman’s rho = 0.65, *p* = 0.01, 95% CI = [0.24 0.88], Figure 3d). All the above correlations were resistant to influences from outliers as the correlations remained significant when eliminating outliers using the Spearman skipped correlation. SA was not significantly correlated with alpha/beta power changes (see supplementary figure S2).

**Figure 3.**
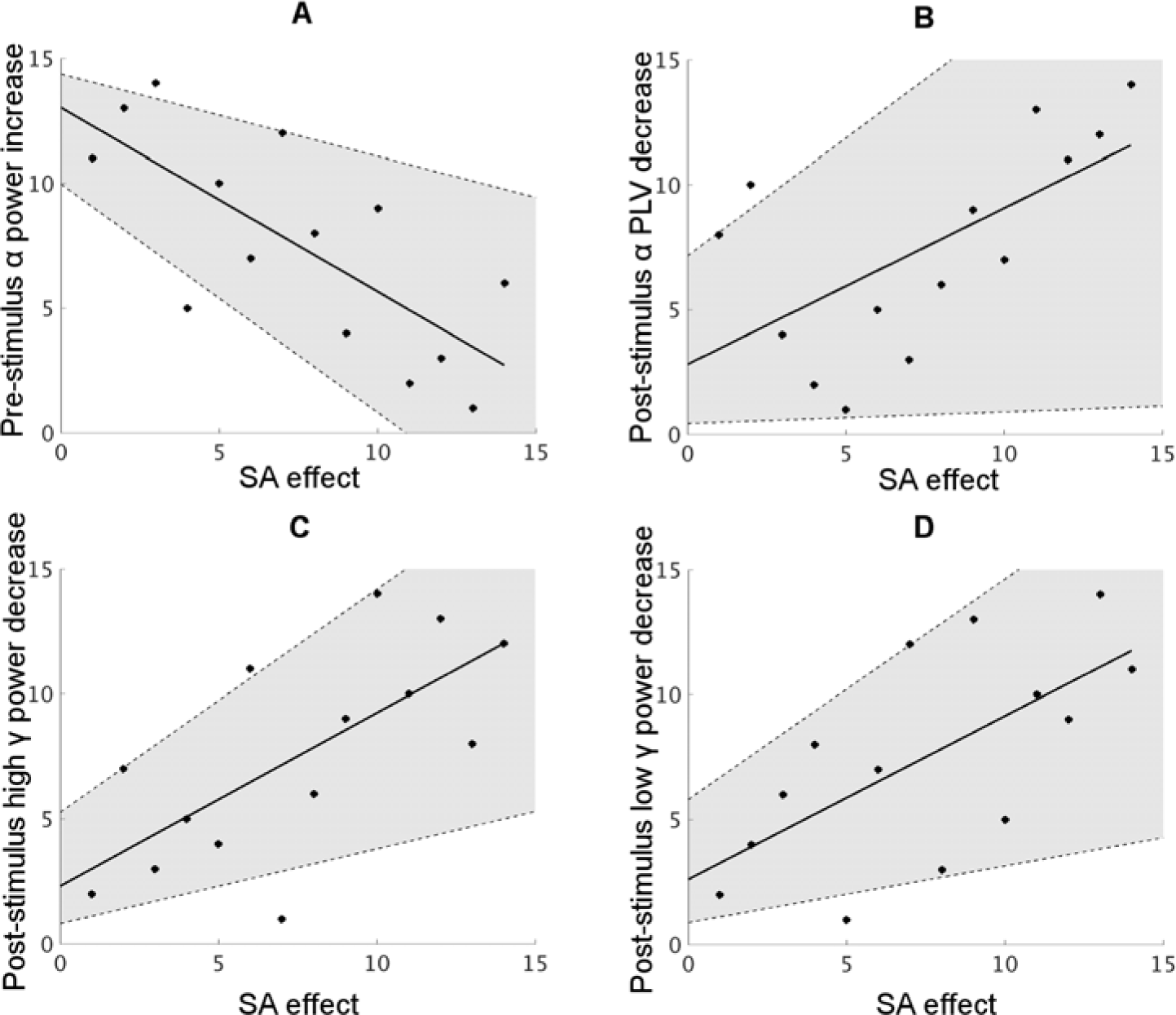
Scatter plots (ranking data) for the correlations between SA in source space and significant power/phase locking changes between conditions. (A) SA is negatively correlated with pre-stimulus alpha power increase (r = −0.74, p = 0.003, CI = [−0.92 −0.33]). (B) SA is positively correlated with post-stimulus alpha phase locking decrease (r = 0.63, p = 0.02, CI = [0.05 0.94]). (C) SA is positively correlated with post-stimulus high gamma power (85-104 Hz) decrease (r = 0.69, p = 0.006, CI = [0.29 0.90]). (D) same with C, but with lower gamma (57-62 Hz) (r = 0.65, p = 0.01, CI = [0.24 0.88]). The solid line indicates a linear fitting to the data points and the shaded area indicates the 95% confidence interval of the correlation.

### Neuronal implementation of sensory attenuation in auditory cortex

Our results so far demonstrate that changes in pre-stimulus alpha power, post-stimulus alpha phase locking and post-stimulus gamma power were most relevant to SA, as evidenced by both, significant between-condition differences and correlations across participants. Because the pre-stimulus low frequency power change (especially in the alpha band) and post-stimulus oscillatory changes are possible candidates for mediating the SA, we investigated if these oscillatory components are correlated amongst each other. Across participants, there was a significant negative correlation between the pre-stimulus alpha power increase *and* the post-stimulus high gamma power decrease (Spearman’s rho = −0.82, p = 0.0003, 95% CI = [−0.94 −0.50]; Figure 4a). Moreover, a significant positive correlation was found between the post-stimulus gamma power decrease and the post-stimulus alpha phase locking decrease (Spearman’s rho = 0.73, p = 0.003, 95% CI = [0.31 0.92]; Figure 4b). Both correlations were resistant to possible influences of outliers (see supplementary material for robust correlation results) and remained significant after Holm-Bonferroni correction. For a full correlation map, see supplementary figure S3.

**Figure 4.**
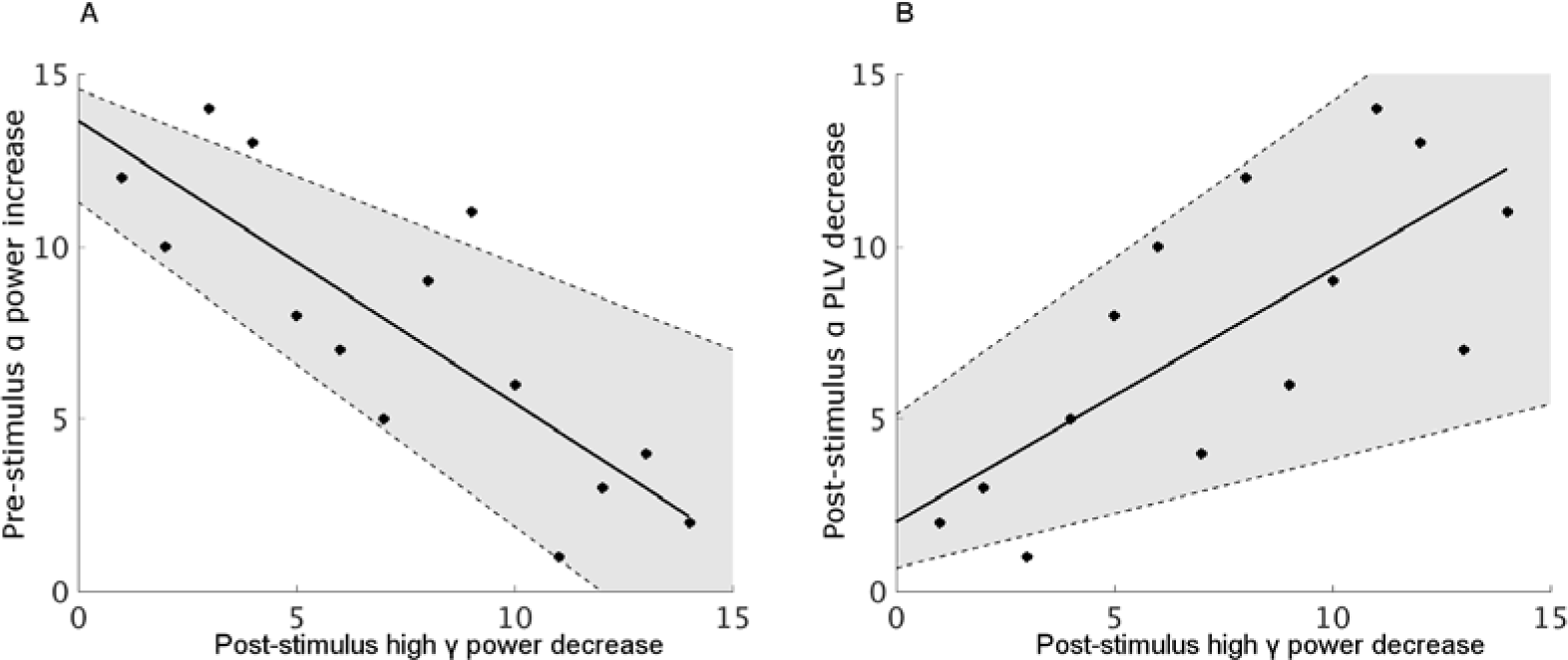
Scatter plots (ranking data) for cross-participant correlations between pre-stimulus alpha power increase and post-stimulus gamma power decrease (A), and between post-stimulus alpha phase locking decrease and post-stimulus gamma power decrease (B). An increase in the pre-stimulus alpha power is associated with a decrease in the post-stimulus gamma power (Spearman’s rho = −0.82, p = 0.0003, 95% CI = [−0.94 −0.50]), and a decrease in the post-stimulus gamma power is associated with a decrease in the post-stimulus alpha phase locking (Spearman’s rho = 0.73, p = 0.003, 95% CI = [0.31 0.92]). The solid line indicates a linear fit to the data points and the shaded area indicates the 95% confidence interval of the correlation.

### Co-variation of the auditory pre-stimulus and post-stimulus oscillatory components related to sensory attenuation in the absence of predictive cues

Next, we tested whether the sequence of events described above (pre-stimulus alpha power relating to post-stimulus gamma power and alpha/beta phase reset) is also present during auditory sensory stimulus processing when no explicit predictions can be formed. Therefore, we tested for the presence of the same correlations within participant in a cross-trial analysis of the passive listening condition in which the inter-trial interval was randomly jittered (between 2000 and 4000 ms). The jittered interval makes the exact onset of the stimulus unpredictable, thus leading to a variation in the participant’s preparedness towards the stimulus.

We correlated pre-stimulus alpha power (8-12 Hz; −300 ms to 0 ms) with post-stimulus gamma power across trials and subjected the individual correlation maps to group statistics. Consistent with our analysis across participants, we found a significant correlation between pre-stimulus alpha power and early post-stimulus gamma power (Figure 5a). The negative sign of the correlation indicated that a high pre-stimulus alpha power was associated with a low post-stimulus gamma power. The pre-stimulus alpha power was also correlated with gamma power (around 85 Hz) starting around 100 ms after stimulus onset.

**Figure 5.**
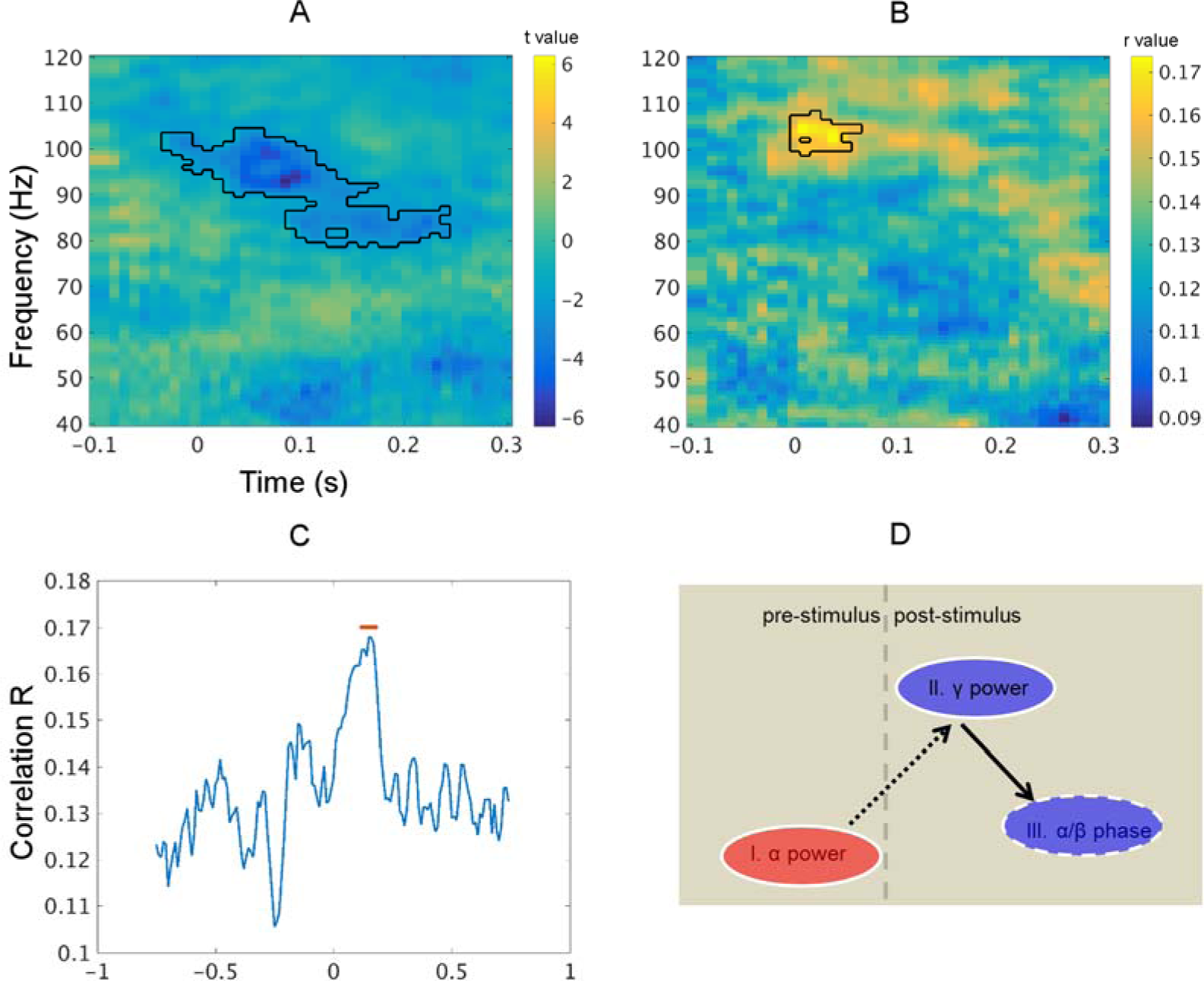
Results from the cross-trial analysis and schematic summary. In the passive jittered condition, pre-stimulus alpha power is correlated with post-stimulus gamma power (A), and post-stimulus alpha/beta phase deviation is correlated with the gamma power at comparable frequency bands and time points (B). Taking the first 40 ms post-stimulus gamma power where significant correlations were found in (B), the temporal dynamics of its correlation with post-stimulus alpha/beta phase deviation is shown in (C). There is a clear peak around 150 ms after the stimulus onset. The red line indicates post-stimulus points where there are significant higher correlations than the baseline period (-750 ms to 0; paired t test without multiple comparison correction; p < 0.05). D is the schematic illustration of the relationship found among oscillatory changes between conditions. The increase of pre-stimulus alpha power is negatively correlated with the decrease of post-stimulus gamma power, which in turn is positively correlated with the decrease of post-stimulus alpha/beta phase locking. This may constitute a sequence of neural information processing (from I to III) which receives further support from the single-trial analysis with the passive jittered condition data. Red colour indicates a relative signal increase in the active condition and blue indicates a decrease. Solid ellipse edge indicates power and dashed line indicates phase. Dashed arrow indicates a negative correlation and solid arrow indicates a positive correlation.

In order to reveal a possible relationship between post-stimulus gamma power and post-stimulus alpha/beta phase across trials (in analogy to the cross participant analysis above), we calculated the phase deviation as the absolute angular difference of a single trial phase to the mean phase across trials for each individual at alpha/beta frequency shortly after stimulus onset. We then correlated this phase deviation with single-trial power across time and frequency. Post-stimulus gamma power was significantly correlated with alpha/beta band phase deviation shortly after stimulus onset (Figure 5b). This correlation was only significant for post-stimulus phase from 11 to 14 Hz (see supplementary Figure S4). Importantly, recalculating the correlation by taking into account only the very early gamma power data (first 40ms of significant post-stimulus correlations) revealed its correlation with alpha/beta phase deviation to peak at a later time point (at around 150 ms, Figure 5c), indicating that gamma power precedes alpha/beta phase resetting.

## Discussion

Although sensory attenuation has been typically studied in the auditory system, the neural mechanisms underlying this phenomenon are likely part of a neural functional architecture that acts along the different stages of sensory processing pathways. In fact, this well-studied effect has been linked to the predictive coding framework that postulates the importance of predictive neural models for general information processing in brain networks (Brown, Adams, Parees, Edwards, & Friston, 2013; Friston, 2005). In this Bayesian framework the brain generates predictions about the environment that are constantly compared to and updated by incoming sensory evidence. The resulting prediction errors are communicated to the next level in the processing hierarchy. An integral part of this theory is the control of gain of these prediction errors that is adjusted according to their expected precision (Friston, Bastos, Pinotsis, & Litvak, 2015). It has been argued that sensory attenuation originates from reduced precision of self-generated sensory information (Brown et al., 2013). Interestingly, brain oscillations provide efficient mechanisms for gain control and are ideal candidates for the neural mechanisms underlying SA.

Our findings support this hypothesis. They replicate the classical SA effect (i.e., self-generated sound elicited a smaller amplitude of M100 component compared to externally generated sound) and show, first, how pre-stimulus changes of auditory alpha band oscillatory power affect auditory stimulus processing and lead to SA. Second, we demonstrate that at the level of single trials SA is reflected in, both reduced broadband power (including gamma) and reduced alpha/beta phase locking when comparing the active and passive periodic condition. Third, we find a significant relationship between pre-stimlus alpha power changes, post-stimulus gamma power changes and post-stimulus alpha/beta phase changes, which may represent a functional sequence of neural information processing steps around the time of stimlus presentation (see Figure 5d). This receives further support from the single trial correlation analysis performed on the passive jittered condition data.

### Pre-stimulus predictors of sensory attenuation

We tested the hypothesis that modulation of low-frequency auditory oscillations are involved in the implementation of sensory attenuation. Indeed, our results demonstrate an enhancement of auditory alpha oscillations (∼10Hz) for the active condition compared to the passive periodic condition before stimulus onset. Low frequency oscillations particularly in the alpha range have previously been related to the suppression of task-irrelevant stimuli or stimulus features (Foxe & Snyder, 2011; Jensen & Mazaheri, 2010) in sensory systems such as visual cortex (Capilla, Schoffelen, Paterson, Thut, & Gross, 2012), auditory cortex (Weisz et al., 2011) and somatosensory cortex (Baumgarten, Schnitzler, & Lange, 2014; Haegens, Händel, & Jensen, 2011).

Therefore the upregulation of alpha oscillations is a viable mechanism for suppression of stimulus evoked activity in auditory cortex when stimulus presentation is self-generated. Indeed, our results are compatible with recent reports of enhanced alpha oscillations for self-uttered sound (Müller et al., 2014) and self-initiated visual stimuli (Stenner et al., 2014). Interestingly, our analysis revealed a significant correlation between alpha power changes and SA, i.e., increased pre-stimulus alpha power changes were associated with increased SA (i.e. more attenuated M100 response) across participants. This provides further evidence for a close relationship between pre-stimulus alpha power modulation and SA and is consistent with reports that pre-stimulus alpha power correlates with early evoked responses (Ploner, Gross, Timmermann, Pollok, & Schnitzler, 2006). This finding is also consistent with results from a recent study that modulation of alpha power reflects the precision of predictions about upcoming stimuli (Bauer et al., 2014). This suggests that individual differences in the ability to predict the sensory consequences of one’s actions are expressed in differences in the modulation of alpha power.

Overall, the significantly increased alpha power for the active compared to the passive periodic condition speaks in favour of an active inhibition of auditory areas at the time of motor preparation as a result of top-down mediated predictions in anticipation of the self-generated (predicted) sensory stimulus.

### Post-stimulus representations of sensory attenuation

The sound evoked response is characterized by an increase of both, oscillatory power and phase locking that is strongest in the theta (4-7 Hz) band (see supplementary figure S1). However, this activity is not modulated between the experimental conditions (active vs passive periodic). Instead, we show here that SA was associated with a significant decrease of power and phase locking in auditory cortex at higher frequencies. This suggests that the mechanisms responsible for SA spare the low-frequency theta component and instead modulate alpha/beta and gamma components. A possible interpretation is that the low-frequency theta component reflects the physical stimulus properties that are unchanged between active and passive periodic conditions whereas higher frequency components reflect more subjective properties of the stimulus that are discussed below in more detail (Gross, Schnitzler, Timmermann, & Ploner, 2007; Iversen, Repp, & Patel, 2009). The fact that SA was associated with a transient broadband power decrease in alpha and beta frequency bands up to almost 40 Hz is in line with a recent study that looked at the top-down modulation of brain responses to simple auditory rhythms (Iversen et al., 2009). Strongest top-down effects were observed in the beta range (while the alpha band was not studied).

Interestingly, high gamma band power was also reduced in the active condition compared to the passive periodic condition, which is consistent with intracranial recordings from patients (Flinker et al., 2010). Animal studies have demonstrated that gamma rhythms are most prominent in supragranular layers suggesting that they mediate feedforward processes (Bastos et al., 2015; Michalareas et al., 2016; van Kerkoerle et al., 2014). In the framework of predictive coding, gamma band activity was suggested to reflect prediction errors for feedforward information processing (Arnal & Giraud, 2012; Bauer et al., 2014; Friston et al., 2015). A predicted stimulus (active condition) is associated with smaller prediction errors thus leading to reduced gamma power.

Another significant difference between active and passive periodic condition shortly after stimulus onset emerged from the phase locking analysis. Alpha/beta phase locking was weaker for active compared to passive periodic condition. This difference is caused by a higher variability of single-trial phase across trials in the active condition. There are different possible explanations for this effect. It could be a byproduct of the reduced alpha/beta post-stimulus power for the active condition. In this scenario the reduced signal-to-noise ratio (SNR) due to the reduced power leads to an artificially reduced (less precise) estimate of single-trial phase. However, this scenario is unlikely for three reasons. First, post-stimulus power is also significantly reduced for high beta and gamma frequency bands without a difference in phase locking. Second, the correlation of post-stimulus gamma modulation to post-stimulus phase locking speaks against a simple SNR-induced effect, especially because post-stimulus gamma modulation correlates with alpha/beta phase but not alpha/beta power (see supplementary figure S3 c&d). Third, the single-trial correlation of gamma power and alpha/beta phase favours a different interpretation where alpha/beta phase is causally linked to gamma amplitude.

### Implications for neural information processing

Both, single-trial analysis and statistical contrasts between conditions revealed a functional neural information processing sequence among pre-stimulus alpha power, post-stimulus gamma power and post-stimulus alpha/beta phase. Prior to the stimulus onset, alpha power controls the gain of local neuronal populations reflecting the precision of the prediction about the incoming stimulus. Mechanistically, this may be implemented by modulating local neuronal excitability levels, known to be indexed by alpha activity (Romei et al., 2008). The pre-stimulus alpha power in the passive jittered condition fluctuated from trial to trial creating differential levels of precision over the incoming stimulus, with low alpha power corresponding to high levels of precision. When the stimulus arrives, any incongruency between the prediction and the actual incoming stimulus (prediction error) is fed forward for further processing through gamma oscillations. Since prediction error is weighted by precision, a negative correlation between pre-stimulus alpha power and post-stimulus gamma power is predicted. This is exactly what we observed (Figure 5a). Next, brain areas from the higher hierarchy processing prediction errors provide feedback to the lower hierarchy through alpha/beta oscillations, which is captured by the significant correlation between post-stimulus gamma power and post-stimulus alpha/beta phase deviation (Figure 5b). The idea of alpha/beta phase acting as top-down signals to resolve the bottom-up prediction error has received support from previous auditory studies (Arnal, Wyart, & Giraud, 2011; Fontolan, Morillon, Liegeois-Chauvel, & Giraud, 2014). In a recent study, Fontolan et al. (2014) showed that gamma power in A1 was modulted by alpha/beta phase in auditory association cortex suggesting the top-down origin of the latter. Our analysis provides further evidence for this by showing that an early gamma power led to a late alpha/beta phase resetting. This temporal asymmetry is an important step for establishing a causal role of gamma power in resetting alpha/beta phase. All these results fit very well with recent findings suggesting that high frequency band oscillations (e.g., gamma) relay feedforward information and that low frequency band (e.g., alpha and beta) oscillations relay feedback information (Bastos et al., 2015; Michalareas et al., 2016; van Kerkoerle et al., 2014).

It is important to note a fundamental difference between the pre- and post-stimulus spectral components. The prestimulus alpha modulation is extended in time and occurs in the absence of a stimulus and likely represents an ongoing oscillation. The post-stimulus effects are partly broadband in nature and are confined to a short time interval where stimulus information is processed. It is therefore unclear to what extent the observed post-stimulus effects originate from changes in brain oscillations. An alternatve explanation is that the two major post-stimulus events (gamma power and alpha/beta phase) reflect a trivial consequence of doing time-frequency analysis in the time window overlapping with evoked responses in the above within- and between-condition analysis. However, this scenario is not consistent with out results.. Specifically, no signficant information was carried in the theta band, where the strongest activation was found in the time window of the evoked componenent after time-frequency analysis. Instead, only the post-stimulus gamma power and alpha/beta phase were identified as critical in the proposed sequence of neural information processing. Furthermore, the frequency bands for interesting events identified in the current study (around 100 Hz for high gamma power and 12-14 Hz for alpha/beta phase) correspond very well with previous studies based on intracranial recordings (Arnal et al., 2011; Flinker et al., 2010). Thus a coherent and consistent explanation of our data requires the existence of functionally distinct frequency components in the recorded brain activity.

## Conclusion

In summary, our results support an involvement of low-frequency auditory oscillations for mediating sensory attenuation. Our findings are consistent with a predictive coding account of sensory attenuation that rests on auditory oscillations for gain control of sensory evidence. They also corroborate recent findings by providing evidence for hierarchical information processing in the brain mediated by gamma (bottom-up) and alpha/beta (top-down) oscillations.

